# Extraordinary physiology of polyphosphate-accumulating *Beggiatoa* mats suggests a key role for phosphate buffering in marine sediments

**DOI:** 10.1101/2024.11.22.624918

**Authors:** Nadezhda Iakovchuk, Jenny Fabian, Olaf Dellwig, Christiane Hassenrück, Heide N. Schulz-Vogt

## Abstract

Filamentous sulfide-oxidizing *Beggiatoa* spp. are widespread in marine coastal environments and can achieve significant biomass because of their substantial size. Their ability to store phosphates in the polymerized form of polyphosphates makes them potentially key players in altering the phosphorus (P) cycle at the sediment-water interface. This study examined phosphate uptake and polyphosphate formation in a P starved culture of the *Beggiatoa* sp. 35Flor strain. Remarkably, even after severe P starvation over five generations, the survival of the cultures was 46%, demonstrating considerable plasticity to different levels of phosphate availability. Under these P-depleted conditions, 23% of filaments still contained polyphosphates, underscoring its critical role in their metabolism. Upon reintroduction of phosphate to starved cultures, an extremely rapid phosphate uptake was observed within the first 10 minutes, with rates reaching up to 298 mmol P g^-1^ protein d^-1^, which is significantly higher than values previously described in the literature for similar-sized organisms. The high phosphate uptake capacity of *Beggiatoa* spp., estimated at 0.6 – 6 mmol m^-2^ d^-1^ for typical densities of filaments in coastal sediments, suggests that these bacteria may play an important role in buffering the phosphate flux in these environments. Thereby, they reduce primary production and subsequent oxygen consumption by other organisms, creating a negative feedback loop that helps maintain ecosystem stability.

**Importance:** Sulfide-oxidizing bacteria of the genus *Beggiatoa* occur ubiquitously in marine coastal sediments and have a large potential to influence phosphate fluxes at the sediment-water interface owing to their ability to accumulate polyphosphate and their large size. However, the extent to which these bacteria can contribute to phosphorus (P) sequestration or release remains poorly assessed. The importance of this study lies in demonstrating the unusual flexibility in adaptation of the *Beggiatoa* sp. 35Flor strain to varying P availability, including extreme P starvation, and its capacity to rapidly uptake and store available phosphate in the form of polyphosphate. When considered at a global scale, these physiological traits could lead to P retention in shallow coastal waters, which, in turn, profoundly impacts ecological stability and ecosystem functioning.

## Introduction

Phosphorus (P) is a vital micronutrient essential for the composition of key biomolecules that govern cellular processes, such as replication, energy transfer and structural integrity; thus, it is often a limiting factor for living organisms (Pasek, 2008). In aquatic environments, P availability is closely linked to iron (Fe) because of the affinity of phosphate for solid Fe oxyhydroxides, and is primarily driven by changes in redox conditions (Föllmi, 1996). Under oxic conditions, phosphate adsorbs onto Fe oxyhydroxides that co-precipitate in the uppermost sediment layer, thereby making phosphate unavailable to most organisms. Conversely, under sulfidic conditions, these Fe oxyhydroxides are reduced thereby releasing bound phosphate back into the porewater. In addition to chemical phosphate binding and release, it is becoming increasingly clear that biological processes themselves such as bacterial polyphosphate (polyP) accumulation have a significant influence on P availability in the system (Gächter & Müller, 2003; Lake et al., 2007). Although many studies have addressed the importance of sediment microbial communities on P dynamics, the significance of polyP-accumulating bacteria remains underestimated (Hupfer et al., 2007; Diaz et al., 2012; Saia et al., 2021).

Polyphosphate is a linear molecule composed of multiple orthophosphate units connected by energy-rich phosphoanhydride bonds (Achbergerová & Nahálka, 2011 and references therein). Prokaryotic organisms synthesize and store polyP in the form of granules, which may be decomposed into orthophosphates or directly utilized for multiple functions (Albi & Serrano, 2016). Polyphopshates have been recognized as an intracellular energy buffer and metabolic regulator; consequently, they have been extensively studied in various prokaryotic organisms (Shiba et al., 1997; Ault-Riché et al., 1998; Kornberg et al., 1999; Rao et al., 2009).

With respect to phosphate availability, three distinguishable conditions that stimulate polyP synthesis in microorganisms have been defined in the literature. Under excess phosphate, luxury uptake occurs through the accumulation of polyP beyond its immediate metabolic needs for future use when phosphate becomes limited (Pauli & Kaitala, 1997; Li & Dittrich, 2019). Conversely, overplus uptake was described when phosphate-deprived cells are re-exposed to phosphate and rapidly accumulate polyP at levels greater than those seen in luxury uptake (Werner et al., 2005). Additionally, a process termed deficiency or starvation response has been recognized for the ability of microorganisms to form polyP, even under low phosphate conditions (Martin et al., 2014; Santoro et al., 2023), although the physiological benefits of this process are not yet fully understood (Dyhrman et al., 2012).

Notably, large sulfur bacteria accumulate polyP extensively due to their size and thus can significantly affect P fluxes in marine sediments, thereby impacting local and global biogeochemical cycles. Such cases have been described for the Namibian shelf, where the species *Thiomargarita namibiensis* was associated with the formation of phosphorites (Schulz & Schulz, 2005; Goldhammer et al., 2010). Among others, the filamentous large sulfur bacteria *Beggiatoa* sp. 35Flor strain was described to cause significant phosphate release under high sulfide fluxes in the absence of oxygen in laboratory experiments (Brock & Schulz-Vogt, 2011), which was later observed in some marine environments (Noffke et al., 2016). Although a systematic estimation of the density of *Beggiatoa* spp. filaments is available only for restricted locations (Jørgensen, 1977; Mußmann et al., 2003; Jørgensen et al., 2010), their presence has been widely reported in coastal marine sediments globally (Strohl & Larkin, 1978; Elliott et al., 2006; Lloyd et al., 2010; Hinck et al., 2011; McKay et al., 2012; Beam et al., 2020; Ravin et al., 2024). Considering their ubiquitous nature, the role of these bacteria in the P cycle is highly under-studied, especially in environments where they do not form a distinct white mat on the sediment surface, but are still present at significant numbers within the sediment. Investigating the response of *Beggiatoa* spp. to different levels of phosphate availability remains a critical gap to understand their potential role in benthic P cycling.

The main objective of this study was to investigate the response of *Beggiatoa* sp. 35Flor to a phosphate overplus and subsequent luxury uptake to evaluate the capacity of bacterial mats to buffer sudden nutrient pulses. We specifically address two main questions: 1. How does a lack of phosphate influence the survival of *Beggiatoa* sp. 35Flor across successive generations? and 2. What are the dynamics of phosphate uptake and the subsequent polyP formation after P reintroduction? To answer these questions, we conducted laboratory experiments in which we subjected the marine *Beggiatoa* sp. 35Flor strain to P starvation for several generations. Subsequently, we introduced an excess of phosphate into the bacterial mat and monitored both phosphate uptake and polyP accumulation. To the best of our knowledge, this is the first attempt to study P starvation and the subsequent polyP dynamics in response to overplus P conditions in *Beggiatoa* species.

## Materials and methods

### Culture and cultivation

The experiment was conducted on filamentous marine sulfide-oxidizing *Beggiatoa* sp. 35Flor strain brought into culture in 2002. It was originally isolated from a single filament of an enriched microbial community from black band disease in scleractinian corals along the coast of Florida. The chemolithoautotrophic *Beggiatoa* sp. strain 35Flor has 6 μm filament thickness and is characterized by distinct polyP and sulfur inclusions (Brock & Schulz-Vogt, 2011). It is accompanied by the heterotrophic and metabolically versatile bacterium *Pseudovibrio* sp. strain FO-BEG1, which is present in low biomass and is required for the growth of this *Beggiatoa* sp. 35Flor under culture conditions (Schwedt, 2011).

The co-culture is maintained in a two-layer mineral semi-liquid agar medium with an opposed gradient of sulfide and oxygen as described by Nelson & Jannasch (1983). It was used as the initial inoculum for growing P-depleted cultures. P-depleted cultures were acquired by cultivating bacteria in modified media without any additional P sources as follows: agar was washed three times with 18.2 MΩ cm water to minimize P contamination and the final agar concentration of the top layer was lowered to 0.2%. The bottom agar layer was supplied with a sulfide concentration of 4 mmol L-1. Because of the high phosphate concentration in the initial stock cultures and the internal storage of polyP in the filaments, it took five generations to acquire cultures in which most of the filaments were free of polyP inclusions confirmed by microscopic observations (see below: Polyphosphate visualization). Each generation was cultivated under controlled conditions at room temperature (20 – 21 °C) in the dark for six days. To create a new generation, a sub-sample (250 – 330 µL) of the bacterial mat was harvested from the previous generation using a sterile Pasteur glass pipette and transferred to freshly prepared medium without additional P source approximately one cm below the air-agar surface interface (Fig. 1).

**Figure 1.**
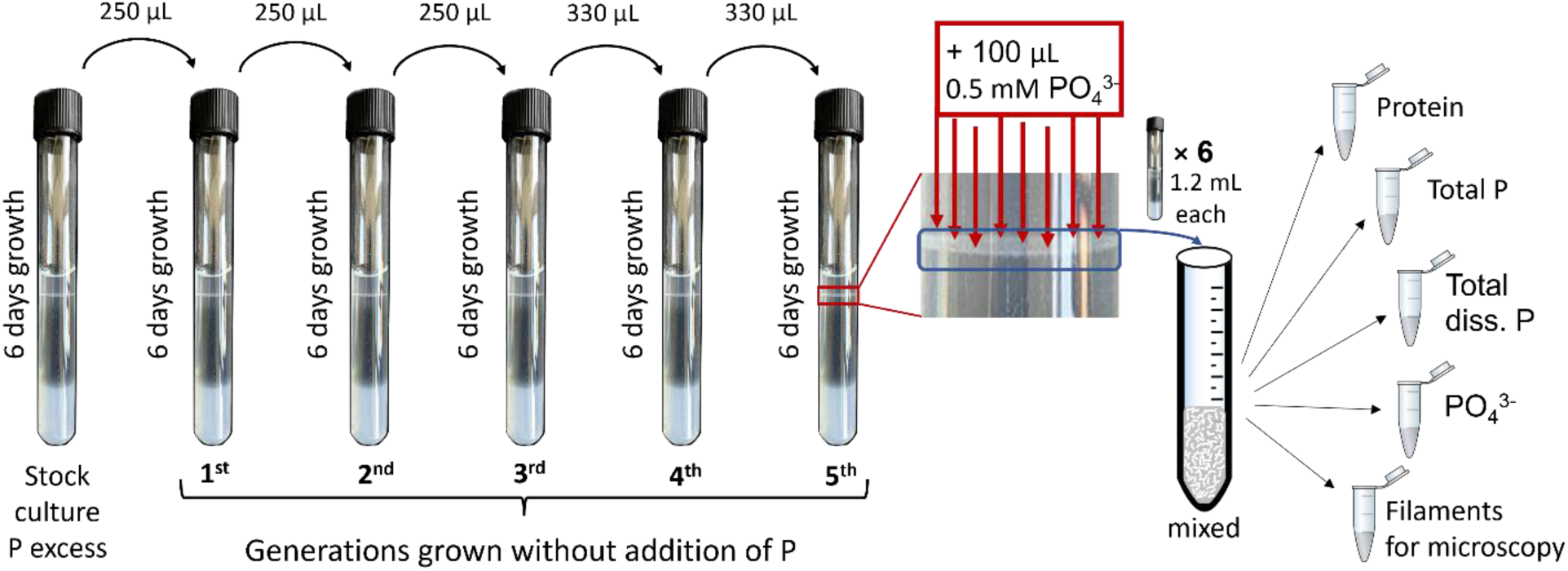
Schematic explaining the experiment design of P-starvation through 5 generations and subsequent incubation with added phosphate. The procedure was repeated for each time point (10 min, 30 min, 2 h, 24h) separately with a new set of cultivation tubes. The entire experiment was repeated 5 times with new culture tubes.

The formation of a distinct white mat 2 – 3 days after inoculation was an indication of successful growth; culture tubes without an established mat were considered as non-vital. Survival for each generation was calculated as the ratio of the number of tubes where a clear bacterial mat was established to the total number of tubes inoculated for a specific generation across all the prepared tubes throughout the pre-experiment investigations.

### Incubation experiment

The experiment began once the fifth generation of P-depleted filaments, grown over six days, was obtained. 100 µL of 0.5 mmol L-1 KH_2_PO_4_ dissolved in artificial seawater were added directly into the mat by randomly placing the tip of the pipet into the mat multiple times (see red arrows in Fig. 1). Throughout the incubation period, samples were collected immediately (within first 10 min, see explanation below) after adding P-supplement, after 30 min, 2 h, and 24 h of incubation. Additionally, we collected P-starved mats, blank samples with sterile medium without P-supplement, and blanks of sterile medium after P addition. Because sampling for dissolved fractions was done by rhizons, it was technically challenging to obtain samples at precise time points. On average, it took 10 min to obtain a sufficient sample volume for further analysis. This time interval was therefore used as a reference point for analyzing the data of samples collected immediately after the addition of phosphate to the bacterial mat. Consequently, all other time points were shifted to the same time interval (10 min) to maintain consistency.

At each sampling time point, bacterial mats from six culture tubes were harvested (1.2 mL each) using a Pasteur glass pipette and combined into a pooled sample in a Falcon tube, which was then homogenized by repeated pipetting up and down (see Fig. 1). To slow down metabolic activity, culture tubes and the pooled sample were submerged in ice during harvesting. The pooled and homogenized sample was then subsampled for the following parameters: total P, protein content, total dissolved P, dissolved inorganic phosphate and filaments for microscopy. For dissolved fractions, samples were taken using rhizons (Rhizosphere Research Products, Wageningen, The Netherlands). The entire experiment was repeated five times over a period of one month to acquire five independent iterations.

As a follow up experiment, we additionally performed a short-term incubation to assess how fast incorporated phosphate appears in the polyP pool of filaments by microscopy. Filaments were collected from the same tube at a two minutes interval over a 30-minute incubation period upon phosphate addition to the fifth generation starved bacterial mat.

### Laboratory measurements

**Total and total dissolved P** were measured by Inductively Coupled Plasma Optical Emission spectroscopy (ICP-OES) at 177.495 nm wavelength with external matrix-matched calibration and online addition of Y as an internal standard. While the total dissolved P fraction was directly measured with ICP-OES after acidification with nitric acid to 2 vol%, the sample material for total P determination was first dried and then digested in closed Teflon vessels at 180°C for 12 h with a mixture of concentrated nitric (2mL) and perchloric (1 mL) acid. After evaporation of the acids, the digestions were fumed-off 3-times with 1 mL 6 M HCl and finally dissolved in 1.5 mL 2 vol% nitric acid. Precision and trueness of the measurement was checked by using the international reference materials SLRS-6 and CASS-6 (NRC) spiked with P and were better than 2.7% and 3.4%, respectively. The total particulate P fraction was calculated by the difference between total and total dissolved P.

Protein content and dissolved inorganic phosphate were both measured on a SPEKOL® 1500 UV VIS Spectrophotometer (Analytik Jena AG, Jena, Germany). **Dissolved inorganic phosphate** was measured using a standard colorimetric method with ascorbic acid (Hansen & Koroleff, 1999).

**Protein content** was determined by the Bradford method (Bradford, 1976) using Bovine Serum Albumin Protein Standard II (Bio-Rad Laboratories, Inc., Hercules, CA, USA). Prior to measurement, protein was extracted from 1 mL sample with 10% TCA (v/v) as described in (Kamp et al., 2008) with slight modifications: after acidic hydrolysis of the agar at 90 °C at 500 rpm, a cooling step was performed on ice for 40 min, and the final pellet was dissolved at 55 °C for 30 min at 250 rpm. Sterile top agar was used as a blank sample.

**Polyphosphates** were visualized by staining filaments with 10 mg L−1 4′,6-Diamidino-2-phenylindole dihydrochloride (DAPI) solution as described previously (Choo et al., 2022). Before staining, 0.5 mL of the sample was fixed with 4% formaldehyde (v/v) and stored at 4 °C for up to 24 h before proceeding with staining. The polyP inclusions were visualized using a fluorescence microscope (Axioskop 2 Mot PLUS, Carl Zeiss, Jena, Germany) with 360/40 nm excitation wavelength. The DAPI-polyP complex exhibits a maximum fluorescence emission of 525 – 550 nm, which results in polyP visualization with a bright yellow-green signal (Gomes et al., 2013).

To determine the proportion of *Beggiatoa* filaments that contained polyphosphate, we counted filaments containing polyP, regardless of the amount and size of granules, and calculated the ratio to the total number of filaments observed on the entire slide.

### Data analysis

All visualization and statistical data analysis was implemented in R (R core team, 2023). To compare the median survival across generations, the non-parametric Kruskal-Wallis test was used, followed by post-hoc Dunn’s test for multiple comparisons using the Benjamini-Hochberg (BH) method using the FSA package (Ogle et al., 2023).

Phosphate uptake rates were calculated assuming a linear behavior between two time points for each iteration of the experiment independently, based on which the mean and standard deviation values for each time period were derived. To express the uptake rate per area, the diameter of the cultivation tube of 1.3 cm was taken into account. The uptake rate per protein was calculated based on the mean value of the protein content for the specific iteration. To convert uptake rate per protein to uptake per biovolume of the filaments, the average conversion coefficient of 39 derived from (Graue, 2007) was used.

Furthermore, an exponential decay model was employed to describe phosphate concentration over time represented by the equation: 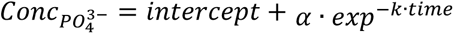. The model was fitted to the data using non-linear least squares regression via the nls() function, with initial parameter estimates set to α = 30, k = 0.5 and intercept = 1. An asymptotic regression was fitted to the particulate P concentrations over time with self-starter function DRC.asymReg() from the aomisc package (Onofri, 2020).

The data is accessible the following: doi.io-warnemuende.de/10.12754/data-2024-0024.

## Results

Survival decreased with each P-starved generation (Fig. 2). A clear drop in survival occurred in the third generation when the median value significantly dropped to 0.75 compared to the first and second generations. By the fifth generation, which was used for the main incubation experiments, the survival had further decreased to 0.46. No significant change was detected in survival between generation three and five.

**Figure 2.**
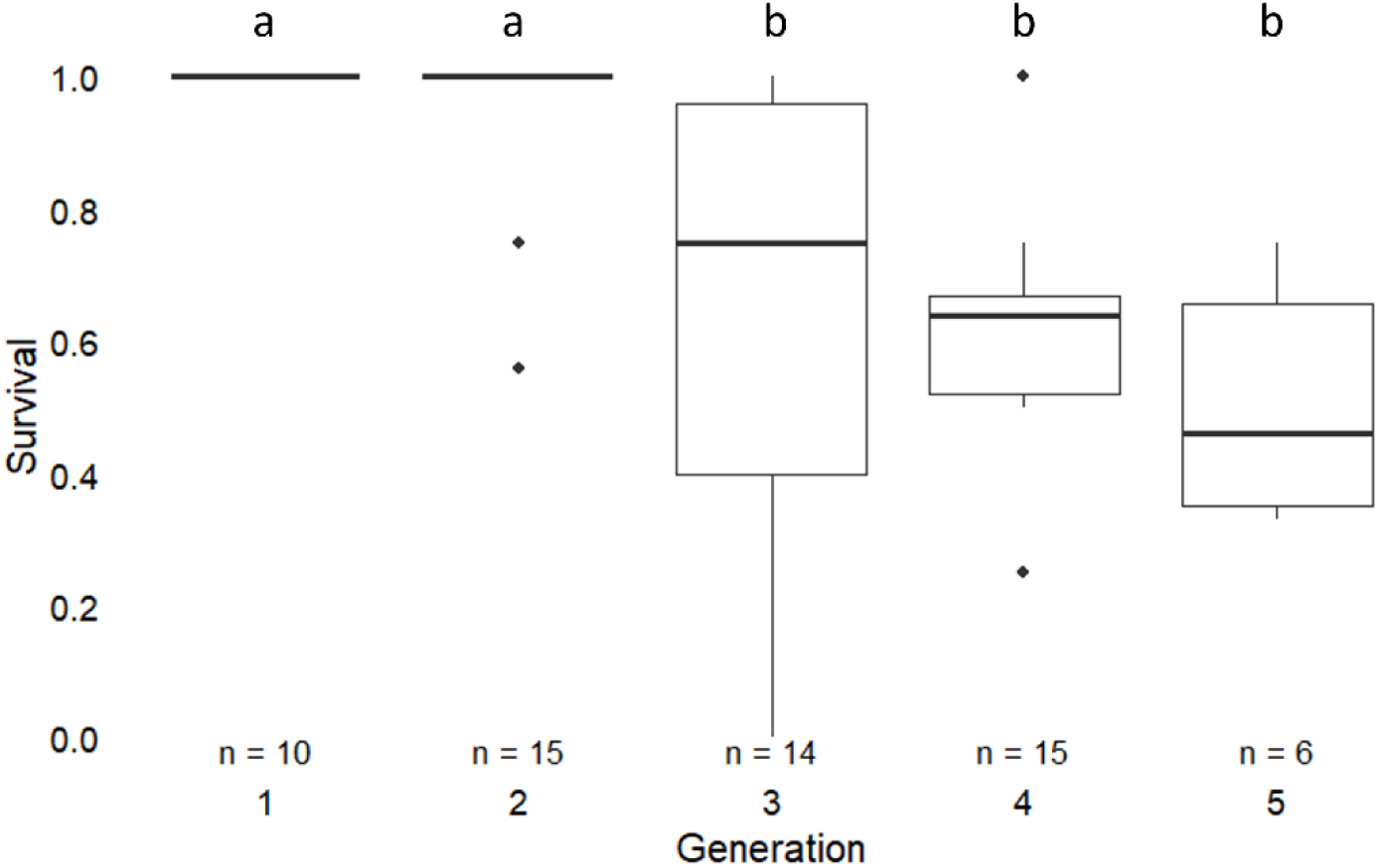
Box-Whisker plots showing survival of P-starved *Beggiatoa* sp. 35Flor cultures over five generations. Boxes show the first and third quartiles; horizontal line: median; whiskers: 1.5 times the interquartile range from the edges of the box; dots represent outliers. Letters (a and b) indicate significant differences (Dunn’s test, BH-adjusted p-value < 0.05), and n indicates the number of independent cultivations with 4 – 40 tubes each.

Fluorescence microscopy of DAPI-stained samples derived from cultures subjected to phosphate starvation for five generations showed that 87% of the filaments did not have any observable polyP inclusions prior to phosphate addition. However, within 30 min upon P-addition, the number of filaments containing at least some polyP increased to 86% and reached 93% by the end of the 24 h of incubation (Fig. 3A). After 30 min of incubation, a clear yellow signal of small round granules was visible, which became larger, emitted a more intense signal after 2 h and further intensified after 24 h of incubation. The polyP distribution varied both across different filaments and within individual filaments. The DAPI-stained filaments (Fig. 3B – E) showed a shift from numerous small granules after 30-minute incubation towards storing polyP in the center of the cell within the main vacuole reaching diameters of up to 4 µm by the end of incubation after 24 h.

**Figure 3.**
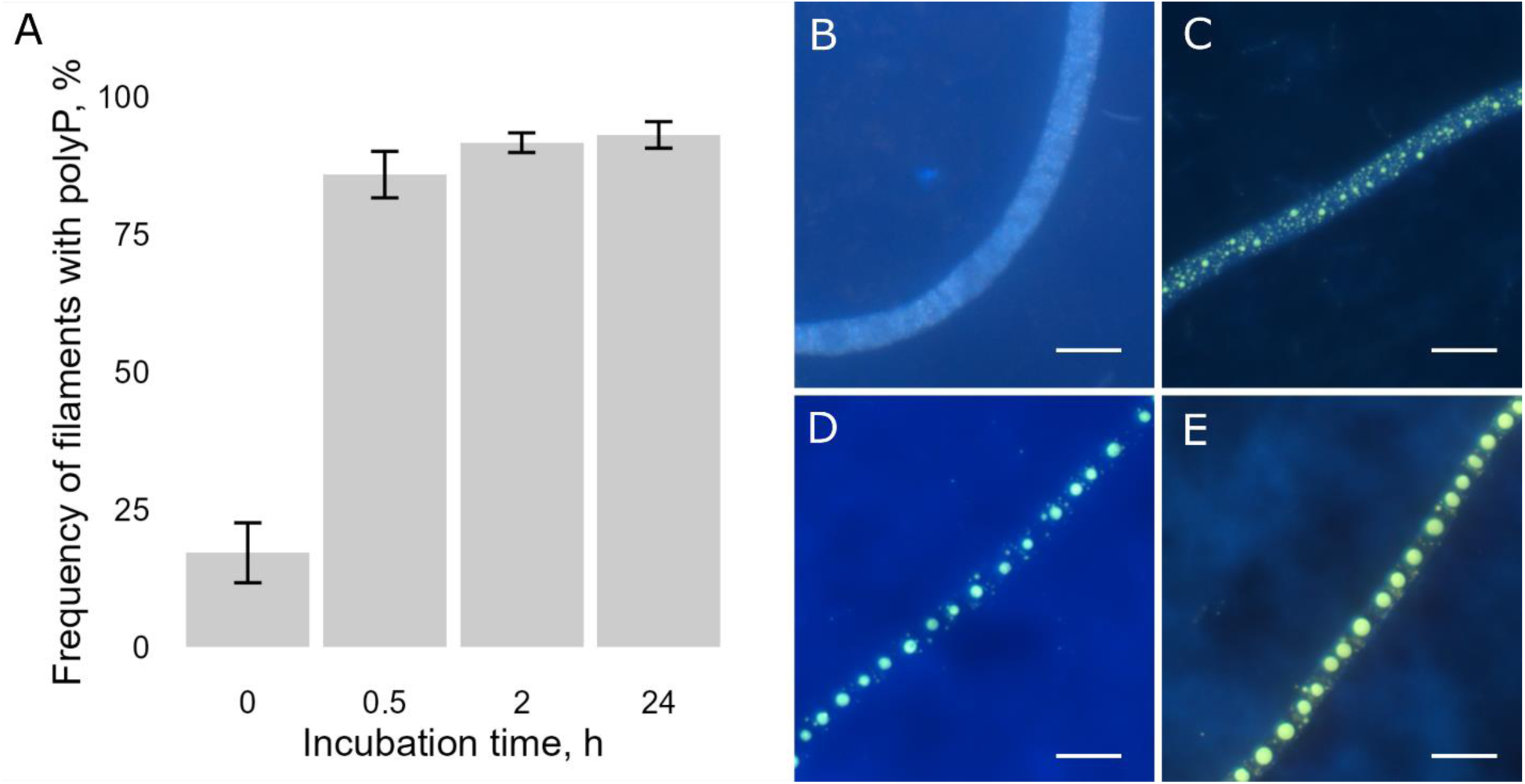
Polyphosphate dynamics in *Beggiatoa* sp. 35Flor following the addition of phosphate to P-starved cultures. A. Percentage of filaments containing polyphosphates (indicated as yes/no) over the course of the experiment; error bars show the standard deviation (n = 5). B – E. DAPI-stained fluorescent images of *Beggiatoa* sp. 35Flor cultures at different times of incubation: B. P-starved culture at the beginning without P-addition, and C. after 30 min, D. after 2 h and E. after 24 h of incubation with P. Bars represent 10 µm.

The average background phosphate concentration in the sterile media, which was used for the starvation cultivations, was 0.2 µmol L^-1^. After 6 days, when the bacterial mat was established in the fifths P-starved generation, the phosphate concentration was below the detection limit for three iterations, for the other two iterations it was 0.11 and 0.12 µmol L^-1^. At the start of the experiment, the addition of KH_2_PO_4_ to the media resulted in an average initial phosphate concentration of 39 µmol L^-1^ measured in sterile media (Fig. 4A). A rapid decrease in the phosphate concentration in the medium was observed within the first 10 min of incubation. This was followed by a nearly 2-fold reduction within the first 30 minutes, and the concentration continued to decrease, reaching an average of 1.78 µmol L^-1^ after 24 hours of incubation. The phosphate concentration decline over time is well described by an exponential decay model (p-value < 0.05 for all coefficients).

**Figure 4.**
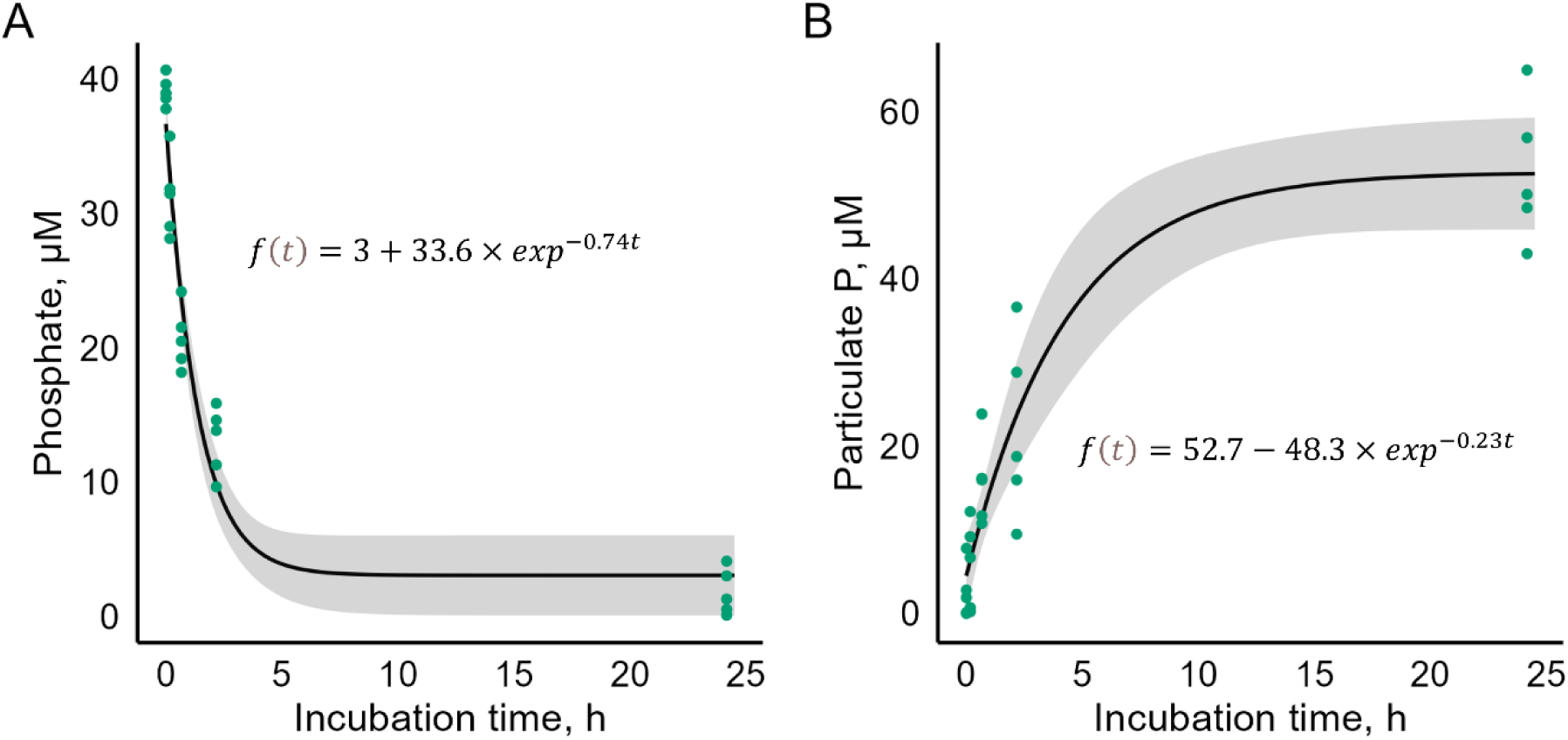
Concentrations of dissolved inorganic phosphate (A) and particulate P (B) over incubation time. Each dot represents individual measurements for five iterations of the experiment. Black line shows the fitted curve, gray ribbon – confidence interval for the fitted curve.

The opposite trend was observed for particulate P (Fig. 4B), which fit to the asymptotic regression (p-value < 0.05 for all coefficients). Rapid accumulation of particulate P occurred within the first 30 min, and the bacterial mat continued to accumulate P until the final sampling point at 24 h of incubation.

Extremely high uptake rates (Table 1) were detected for the time interval between 0 and 10 min, followed by a gradual decrease throughout the rest of the incubation period.

**Table 1.**
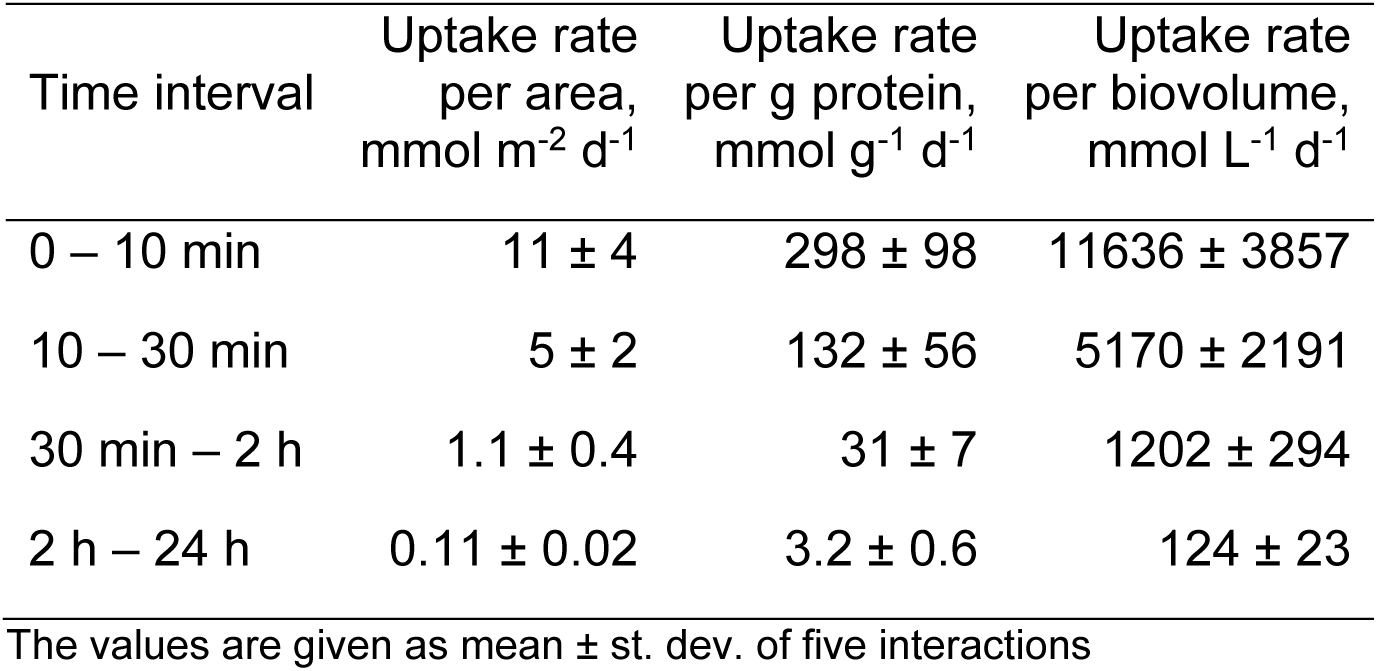
Mean phosphate uptake rates of *Beggiatoa* sp. 35Flor at different time intervals assuming linear relationship.

| Time interval | Uptake rate<br>per area,<br>mmol m<sup>-2</sup> d<sup>-1</sup> | Uptake rate<br>per g protein,<br>mmol g<sup>-1</sup> d<sup>-1</sup> | Uptake rate<br>per biovolume,<br>mmol L<sup>-1</sup> d<sup>-1</sup> |
| --- | --- | --- | --- |
| 0 – 10 min | 11 ± 4 | 298 ± 98 | 11636 ± 3857 |
| 10 – 30 min | 5 ± 2 | 132 ± 56 | 5170 ± 2191 |
| 30 min – 2 h | 1.1 ± 0.4 | 31 ± 7 | 1202 ± 294 |
| 2 h – 24 h | 0.11 ± 0.02 | 3.2 ± 0.6 | 124 ± 23 |

The values are given as mean ± st. dev. of five interactions

## Discussion

### Survival at different generation of P-starved cultures

Healthy *Beggiatoa* populations, in the nature as well as in the culture conditions, aggregate in mats located at the oxic-anoxic interface, where they thrive (Nelson & Jannasch, 1983; Mußmann et al., 2003). They do so to gain access to both sulfide, which acts as electron donor, and oxygen to generate energy for their metabolism (Nelson & Jannasch, 1983). This behavior is driven by negative oxygen taxis and was observed even in the absence of sulfide (Møller et al., 1985). In our experiment, tubes considered non-vital, in most cases, still contained motile individual filaments, but these filaments fail to aggregate at the oxygen-sulfide interface and did not show increase in biomass in the later phases for reasons that are not fully understood. A similar distribution pattern of filaments was observed by Dunker et al. (2011) under conditions when the sulfide and oxygen zones did not overlap. This caused filaments to glide longer distances while using internally stored nitrate as alternative electron acceptors. It is possible that the urgent need for P overrides the typical chemotactic response of filaments to oxygen, causing them to glide beyond the oxygen-sulfide interface to scavenge for available phosphate, which could be present due to background concentration in the media. On the one hand, cells may die because of the inability to generate sufficient energy to support metabolism caused by a mismatch of electron donors and acceptors. On the other hand, gliding of filaments into oxygenated parts of the media may lead to cell death due to their inability to mitigate oxidative stress, as *Beggiatoa* spp. typically lack the catalase enzyme critical for oxidative stress response (Larkin & Strohl, 1983).

An alternative explanation for the inability to form a mat could be attributed to the reduced gliding speed resulting from the missing polyP energy source. Such link between polyP and swimming motility was postulated for pathogenic bacteria (Rashid & Kornberg, 2000), where the authors observed reduced motility due to absence of polyP in mutant strains. A similar connection has also been hypothesized for the polyP-accumulating chemolithoautotrophic sulfur bacteria *Sulfurimonas* subgroup GD17, inhabiting the pelagic redox zone of the Baltic Sea (Möller et al., 2019). The authors linked the energy released from the degradation of polyP to the energy required for the movement of the flagella. Consequently, polyP or P starvation in the present study can induce a decreased ability to find the optimal position across the oxygen-sulfide gradient in the media to sustain growth.

A significant drop in the number of tubes successfully forming a mat after three generations of P-starved cultures (Fig. 2) indicates that P depletion directly affects the survival of *Beggiatoa* sp. 35Flor. This finding aligns with the importance of P as a critical element for many biological molecules, being essential for structural, metabolic, and regulatory functions (Pasek, 2008). Bacteria have a very low cell quota for P in general and can survive at the traces of phosphate in the environment (Godwin & Cotner, 2015). Nevertheless, P starvation induces various changes, including reduced growth, morphological alterations and various physiological adaptations (Seeger & Jerez, 1993; Romano et al., 2015). Poor survival due to P starvation has been reported for *E. coli* (Himeoka et al., 2022), *Pseudomonas putida* (Eberl et al., 1996) and other bacteria (Rao et al., 2009 and references therein). Our data show evidence that long-term P-depletion causes a significant decline in culture survival of *Beggiatoa* sp. 35Flor.

### Crucial role of PolyP in survival

We decided to perform the experiment with the fifth generation of P-starved cultures because only in this generation, we could clearly document the absence of polyP in the majority (87%) of the filaments (Fig. 3A). No additional P was added to the system, only traces of P from different media components (mostly agar) were available (mean 0.2 µmol L^-1^) for the bacteria. It was surprising to observe that *Beggiatoa* sp. 35Flor filaments were able to grow for at least up to five generations under these extreme P-depleted conditions, indicating their extreme resilience to different phosphate levels.

Despite this, even in the fifth generation P-starved cultures, approx. 23% of the *Beggiatoa* sp. 35Flor population still contained polyP (Fig. 3A). Presence of polyP inclusions even at such extreme P-depleted conditions could indicate that polyP not only acts as a reservoir of inorganic phosphate for other molecules (for example, ATP and phospholipids), but may also play an important role in more specific metabolic processes related to critical survival mechanisms. Previous research demonstrated that under various stress conditions, polyP is linked to a general stress response by inducing the expression of rpoS genes responsible for transcription regulating sigma factors (Shiba et al., 1997). This observation was confirmed for *Beggiatoa* sp. 35Flor strain by Langer et al. (2018). The authors showed that the polyP turnover rate in this strain remains high under both favorable (oxic, low sulfide flux) and stressful conditions (anoxic, high sulfide flux), underlining the importance of polyP in stress response and subsequent survival metabolism. Based on the decline in survival of filaments in generations, we can assume that long-term P starvation, to which *Beggiatoa* sp. 35Flor was exposed to in our study, caused stressful growth conditions stimulating a stress response. Furthermore, we hypothesized that the absence of polyP due to starvation could potentially suppress metabolic survival mechanisms. In this respect, the 23% of filaments that were able to maintain polyP in their cells possess an advantage over the rest of the filaments and might play a crucial role in bacterial mat establishment during cultivation.

### Overplus rapid phosphate uptake rate

After adding phosphate to the starved cultures, we observed extremely high phosphate uptake rates by the marine sulfur bacterium *Beggiatoa* sp. 35Flor. In general, nutrient uptake in bacteria can be mathematically described by the Michaelis-Menten kinetic equation, which considers enzyme-catalyzed reactions of active transport. This model assumes the rates to be dependent on the initial substrate concentration and is limited by the number of enzymes present in the cell membrane (Button, 1985). As such, the theoretical maximum phosphate uptake rate of 35. 7 mmol g^-1^ protein d^-1^ under overplus phosphate was reported for the typical bacteria of activated sludge from wastewater treatment plants, *Acinetobacter* spp. (Pauli & Kaitala, 1997). For *Beggiatoa* sp. 35Flor mats, we calculated a phosphate uptake rate of 298 mmol g^-1^ protein d^-1^ (Table 1) over the first 10 min of incubation. This rate is more than eight times higher than reported for *Acinetobacter* spp. This discrepancy could be attributed to the larger cell volume of *Beggiatoa*, which allows greater polyP storage. A reasonable comparison for *Beggiatoa* sp. 35Flor would be with filamentous bacteria of similar sizes, such as freshwater cyanobacteria *Oscillatoria agardhii*, which has been shown to form polyP under overplus conditions (Konopka et al., 1993). The phosphate uptake rate for *O. agardhii* under P-depleted conditions was approximately 54 mmol g^-1^ protein d^-1^ (Riegman & Mur, 1984), and thus nearly six times lower than for *Beggiatoa* sp. 35Flor. Given the comparable sizes of their filaments, other factors may explain the significant differences in uptake rates. One example could be the duration of P starvation and differences in the considered low P concentrations in the experimental design. Another possibility to explain such variations could lay in the differences in physiology and the genetic make-up of different bacterial species.

The rapid uptake of phosphate observed during the initial 10 min of incubation agrees with the typical “overplus” response observed in starved bacteria. In phosphate-deprived environments, most bacteria induce the high-affinity phosphate transport system PstSCAB, which actively transports phosphate, even in environment with limited phosphate availability (Martín & Liras, 2021). After phosphate is re-introduced, microorganisms switch from the high-affinity PstSCAB to the low-affinity PitH phosphate transport system (Martín & Liras, 2021). This switch is not instantaneous, leading to fast initial uptake rates. This phenomenon has been well-documented in cyanobacteria (Jansson, 1993; Solovchenko et al., 2020), various heterotrophic bacteria (Medveczky & Rosenberg, 1971; Van Veen et al., 1993; Gebhard et al., 2006), and yeast (Werner et al., 2005). Because our first sample was collected at maximum 10 min after phosphate addition, the potential effect of the high-affinity transport system may have been underestimated, as previous studies have shown that this process can occur within a few minutes (Jansson, 1993). This finding is also supported by our observations of polyP formation during short-term incubation. Notably, the polyP began to appear after only two minutes of incubation with phosphate (Fig. S1), which is consistent with observations made for *Nostock* by Solovchenko et al. (2020). They postulated that such rapid polyP transport into vacuoles serves as a safety mechanism to prevent the accumulation of short-chain polyP due to its potential toxicity coming from scavenging divalent cations. Indeed, this effect has been documented in yeast cultures, and the incorporation of polyP into acidocalcisomes was proposed as a protective mechanism (Gerasimaite et al., 2014). This assumption is consistent with the observation for *Beggiatoa* sp. 35Flor, where polyP was demonstrated to be stored in acidocalcisome-like vacuoles (Brock et al., 2012).

### Phosphate buffering capacity of *Beggiatoa* spp

Although we could not directly measure polyP, and therefore cannot conclusively state that all the assimilated phosphate was converted into polyP, our microscopic observations (Fig. 3B – E) strongly support the hypothesis of phosphate storage in the form of polyP. We calculated that 100% of added phosphate was incorporated into the particulate pool within 24 hours of incubation (Suppl. Material). Since we did not detect a notable increase in the protein content of the samples, we could therefore assume that the majority of consumed phosphate was stored as polyP. The fact that *Beggiatoa* sp. 35Flor stores all assimilated phosphate in form of polyP demonstrates their high buffering capacity for fluctuating phosphate concentrations in the environment.

To estimate the possible effect of *Beggiatoa* spp. mat on fluctuating phosphate concentrations under environmental conditions, we used the phosphate uptake starting at 30 minutes of incubation, which can be presumably described as ’luxury uptake.’ This is supported by the observation that the filaments already contained polyP granules (Fig. 3B – E) to the extent similar to the growing conditions with P excess and are thus not substantially phosphate-deprived. This phosphate luxury uptake ranges between 124 (2 – 24 h) and 1202 (30 min – 2 h) mmol L^-1^ d^-1^ (Table 1). To extrapolate the obtained phosphate uptake to in-situ conditions, the density of filaments in natural habitats should be considered. The biovolume of *Beggiatoa* spp. filaments has been estimated in only a few published studies, as summarized in Table 2. These studies cover a range of environments, including the Peruvian upwelling system, intertidal mud flats, and the brackish sediments of Limfjorden. However, they may serve as representatives of similar environments globally. The Peruvian upwelling system, for example, represents large portions of the continental west coasts (e.g., California, Chile, Namibia), while intertidal mud flats are prevalent in numerous coastal ecosystems (Murray et al., 2019). Similarly, the brackish sediments of Limfjorden share characteristics with the Baltic Sea and other brackish environments. Thus, the listed habitats, although specific, are representative of broader coastal environments, suggesting that the biovolume data could potentially be applicable to other regions globally.

**Table 2.** Overview of *Beggiatoa* spp. distribution, biovolume, and filament diameter across different locations.

| Location | Environment | Biovolume, $\text{cm}^3 \text{m}^{-2}$ | Filament diameter range (dominant), $\mu\text{m}$ | Examined depth of the sediment | Reference |
| --- | --- | --- | --- | --- | --- |
| Limfjorden, Denmark | Brackish sediment, 4 - 12 m water depth | 5 – 20 | 3 – 35 (15) | depth integrated up to 5 cm | Jørgensen, 1977 |
| Limfjorden, Denmark | Brackish sediment, 4 - 12 m water depth | 14 – 16 | 3 – 40 | depth integrated up to 5 cm | Mußmann et al., 2003 |
| Wadden Sea (Dangast/Jadebay) | intertidal mud flat | 0.6 – 1 | 9 - 12 | depth integrated up to 5 cm | Mußmann et al., 2003 |
| Svalbard | Fjord sediments | 1.13 – 3.36 | 2 – 50 (2 – 5) | depth integrated up to 5 cm | Jørgensen et al., 2010 |
| Peruvian shelf | upwelling system | 0.1 – 0.7 | 5 – 40 | upper 1 cm surface sediment | Langer, 2019 |

Given that the biovolume data discussed previously could be extrapolated on broader regions, we can now apply these values to estimate the phosphate uptake by *Beggiatoa* spp. in natural habitats. Assuming conservatively an average *Beggiatoa* spp. biovolume of 5 cm^3^ m^-2^ across the locations listed in Table 2 and applying the range of phosphate uptake rates per biovolume of filaments calculated in this study (124 – 1202 mmol L^-1^ d^-1^), the average phosphate uptake of such natural environments would be approximately 0.6 – 6 mmol m^-2^ d^-1^. These values are of the same order of magnitude as the phosphate uptake rate of 1.2 mmol m^-2^ d^-1^ detected in Namibian sediments under oxic conditions, where the activity of large sulfur bacteria was studied using a ³³P-radiotracer in laboratory incubations (Goldhammer et al., 2010). The authors linked bacterial phosphate uptake to apatite formation and postulated the importance of sulfide-oxidizing bacteria as P sinks. Similarly, an earlier study from the same location linked phosphorite deposits to the activity of *Thiomargarita namibiensis*, indicating that large sulfur bacteria are crucial in driving phosphorite formation under anoxic conditions (Schulz & Schulz, 2005).

So far, *Beggiatoa* spp. and other large sulfur bacteria have been primarily associated with phosphate release under anoxic and highly sulfidic conditions (Brock & Schulz-Vogt, 2011). However, less attention has been paid to their additional functional roles related to the P cycle in ecosystems where the formation of P-associated minerals does not occur. *Beggiatoa* mats have been reported from various locations, including the shallow coastal waters of Denmark and Germany (Jørgensen, 1977; Mußmann et al., 2003), intertidal flats (Mußmann et al., 2003; Beam et al., 2020), coastal zones of Chile (Aranda et al., 2015), mangrove sediments (Jean et al., 2015), Arctic ocean (Jørgensen et al., 2010; Ravin et al., 2024), deep sea (Lloyd et al., 2010; McKay et al., 2012) as well as freshwater habitats (Strohl & Larkin, 1978) and hypersaline lakes (Hinck et al., 2011). However, *Beggiatoa* spp. do not always form consistently noticeable white mats at the sediment surface and could thus be easily overlooked. Routine investigations of sediment layers have revealed that sometimes the peak of *Beggiatoa* spp. biomass occurs below the sediment surface (Jørgensen, 1977; Mußmann et al., 2003), potentially leading to an underestimation of these groups on a larger spatial scale.

Since marine sediments could harbor similar biovolumes of *Beggiatoa* spp. as those reported in Table 2, we hypothesize that the phosphate uptake rate and polyP storage capacity of *Beggiatoa* spp. are sufficient to quantitatively affect phosphate release into bottom waters coming from diffusive porewater fluxes from deeper parts of the sediment, similar to what have been shown for cyanobacterial mats acting as P-sponge at the sediment surface (Choo et al., 2022). This phosphate sequestration and retention in sediments by *Beggiatoa* spp. helps to mitigate effects of eutrophication caused by release of nutrients in the water column. Thus, apart from preventing sulfide from diffusing out of the sediment, thriving *Beggiatoa* mats can also regulate oxygen availability and mitigate the spread of oxygen deficiency in aquatic ecosystems. On the downside, *Beggiatoa* mats confronted with toxically high levels of sulfide and anoxia will start to release formally stored phosphate in large quantities (Brock et al., 2011), thereby abruptly reversing their role from a mediator to a promotor of eutrophication. These processes highlight the shifting dynamics of the polyP pool, a crucial yet poorly understood phosphorus reservoir in natural ecosystems.

**In conclusion**, our study demonstrates that *Beggiatoa* sp. strain 35Flor can endure severe P depletion, likely facilitated by polyP storage, which appears to play a crucial role in its survival under P-starved conditions. We also observed exceptionally rapid phosphate uptake rates under P-overplus reintroduction, which highlights the unique physiological capabilities of this bacterium. While further studies involving environmental samples are needed to confirm the role of *Beggiatoa* spp. mats in buffering phosphate fluxes, our findings suggest that phosphate sequestration by *Beggiatoa* spp. could significantly mitigate nutrient bioavailability to other organisms. The potential capability of *Beggiatoa* mats to maintain ecosystem stability might give them a crucial role in deciding the faith of benthic environments faced with eutrophication and spreading anoxia.

## Acknowledgments

We gratefully acknowledge Christin Laudan for her assistance with culture cultivation and additional laboratory support. Our sincere thanks go to the members of the Baltic TRANSCOAST Research Training Group for their invaluable support and advice. We also appreciate Simeon Choo for his initial discussions and guidance on phosphate measurements and Jan Henkel for giving valuable feedback on the manuscript.

This study was funded by Deutsche Forschungsgemeinschaft (DFG) under grant number GRK 2000. This is Baltic TRANSCOAST publication number GRK2000/00xx (will be provided after acceptance).

## Supplementary materials

Assuming the theoretical maximum of P in the incubation experiment as the sum of supplied P in the form of phosphate of 0.05 µmol P (when calculated from 100 µL × 0.5 mmol stock solution added to the mat) and the initial P content of the starved bacterial mat of 0.006 µmol P (5.0 µmol L^-1^ × 1.2 mL sample taken), the available P during the incubation would be 0.056 µmol. At the 24 h incubation time point, the mean particulate P was 52.7 µmol L^-1^, which is equivalent to 0.0632 µmol of P (52.7 µmol L^-1^ × 1.2 mL sample taken). This value is slightly higher (about 12%) than the assumed theoretical maximum of P in the incubation. However, considering the analytical error of the measurement, this would indicate 100% incorporation of added P into the particulate pool. Since we did not detect a notable increase in the protein content of the samples, we could therefore assume that the majority of consumed phosphate was stored as polyP.

---

**Figure S1.**
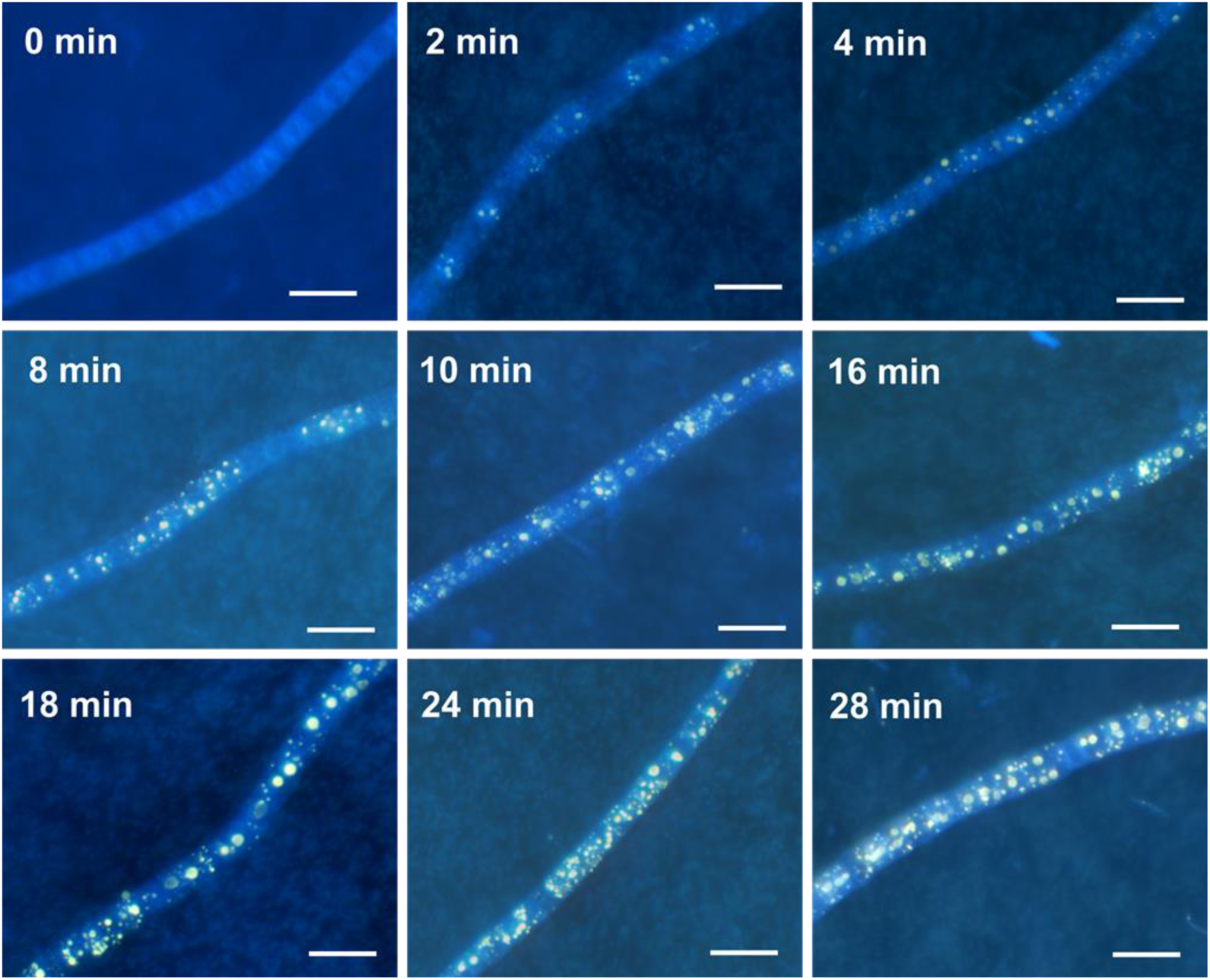
DAPI-stained fluorescent images of *Beggiatoa* sp. 35Flor cultures from short-term incubation experiment with 2 min sampling intervals. 0 min – P starved filaments before addition of phosphate. Bar is 10 µm.

## References

1. Achbergerová, L., & Nahálka, J. (2011). Polyphosphate - an ancient energy source and active metabolic regulator. In Microbial Cell Factories (Vol. 10). 10.1186/1475-2859-10-63

2. Albi, T., & Serrano, A. (2016). Inorganic polyphosphate in the microbial world. Emerging roles for a multifaceted biopolymer. World Journal of Microbiology and Biotechnology, 32(2), 1–12. 10.1007/s11274-015-1983-2

3. Aranda, C. P., Valenzuela, C., Matamala, Y., Godoy, F. A., & Aranda, N. (2015). Sulphur-cycling bacteria and ciliated protozoans in a Beggiatoaceae mat covering organically enriched sediments beneath a salmon farm in a southern Chilean fjord. Marine Pollution Bulletin, 100(1), 270–278. 10.1016/j.marpolbul.2015.08.040

4. Ault-Riché, D., Fraley, C. D., Tzeng, C.-M., & Kornberg, A. (1998). Novel Assay Reveals Multiple Pathways Regulating Stress-Induced Accumulations of Inorganic Polyphosphate in *Escherichia coli*. In Journal of Bacteriology (Vol. 180, Issue 7).

5. Beam, J. P., George, S., Record, N. R., Countway, P. D., Johnston, D. T., Girguis, P. R., & Emerson, D. (2020). Mud, Microbes, and Macrofauna: Seasonal Dynamics of the Iron Biogeochemical Cycle in an Intertidal Mudflat. Frontiers in Marine Science, 7. 10.3389/fmars.2020.562617

6. Bradford, M. M. (1976). A Rapid and Sensitive Method for the Quantitation of Microgram Quantities of Protein Utilizing the Principle of Protein-Dye Binding. Analytical Biochemistry, 72, 248–254. 10.1016/j.cj.2017.04.003

7. Brock, J., Rhiel, E., Beutler, M., Salman, V., & Schulz-Vogt, H. N. (2012). Unusual polyphosphate inclusions observed in a marine *Beggiatoa* strain. International Journal of General and Molecular Microbiology, 101(2), 347–357. 10.1007/s10482-011-9640-8

8. Brock, J., & Schulz-Vogt, H. N. (2011). Sulfide induces phosphate release from polyphosphate in cultures of a marine *Beggiatoa* strain. ISME Journal, 5(3), 497–506. 10.1038/ismej.2010.135

9. Button, D. K. (1985). Kinetics of Nutrient-Limited Transport and Microbial Growth. Microbiological Rewiers, 49(3), 270–297. https://journals.asm.org/journal/mr

10. Choo, S., Dellwig, O., Wäge-Recchioni, J., & Schulz-Vogt, H. N. (2022). Microbial-driven impact on aquatic phosphate fluxes in a coastal peatland. Marine Ecology Progress Series, 702, 19–38. 10.3354/meps14210

11. Diaz, J. M., Ingall, E. D., Snow, S. D., Benitez-Nelson, C. R., Taillefert, M., & Brandes, J. A. (2012). Potential role of inorganic polyphosphate in the cycling of phosphorus within the hypoxic water column of Effingham Inlet, British Columbia. Global Biogeochemical Cycles, 26(2). 10.1029/2011GB004226

12. Dunker, R., Røy, H., Kamp, A., & Jørgensen, B. B. (2011). Motility patterns of fillamentous sulfur bacteria, *Beggiatoa* spp. FEMS Microbiology Ecology, 77(1), 176–185. 10.1111/j.1574-6941.2011.01099.x

13. Dyhrman, S. T., Jenkins, B. D., Rynearson, T. A., Saito, M. A., Mercier, M. L., Alexander, H., Whitney, L. A. P., Drzewianowski, A., Bulygin, V. V., Bertrand, E. M., Wu, Z., Benitez-Nelson, C., & Heithoff, A. (2012). The transcriptome and proteome of the diatom *Thalassiosira pseudonana* reveal a diverse phosphorus stress response. PLoS ONE, 7(3). 10.1371/journal.pone.0033768

14. Eberl, L., Givskov, M., Sternberg, C., Marller, S., Christiansen2, G., & Molin’, S. (1996). Physiological responses of *Pseudornonas putida* KT2442 to phosphate starvation. Microbiology, 142, 155–163.

15. Elliott, J. K., Spear, E., & Wyllie-Echeverria, S. (2006). Mats of *Beggiatoa* bacteria reveal that organic pollution from lumber mills inhibits growth of *Zostera marina*. Marine Ecology, 27(4), 372–380. 10.1111/j.1439-0485.2006.00100.x

16. Föllmi, K. B. (1996). The phosphorus cycle, phosphogenesis and marine phosphate-rich deposits. Earth-Science Reviews, 40, 55–124.

17. Gächter, R., & Müller, B. (2003). Why the phosphorus retention of lakes does not necessarily depend on the oxygen supply to their sediment surface. Limnology and Oceanography, 48(2), 929–933. 10.4319/lo.2003.48.2.0929

18. Gebhard, S., Tran, S. L., & Cook, G. M. (2006). The Phn system of *Mycobacterium smegmatis*: A second high-affinity ABC-transporter for phosphate. Microbiology, 152(11), 3453–3465. 10.1099/mic.0.29201-0

19. Gerasimaite, R., Sharma, S., Desfougères, Y., Schmidt, A., & Mayer, A. (2014). Coupled synthesis and translocation restrains polyphosphate to acidocalcisome-like vacuoles and prevents its toxicity. Journal of Cell Science, 127(23), 5093–5104. 10.1242/jcs.159772

20. Godwin, C. M., & Cotner, J. B. (2015). Stoichiometric flexibility in diverse aquatic heterotrophic bacteria is coupled to differences in cellular phosphorus quotas. Frontiers in Microbiology, 6. 10.3389/fmicb.2015.00159

21. Goldhammer, T., Brüchert, V., Ferdelman, T. G., & Zabel, M. (2010). Microbial sequestration of phosphorus in anoxic upwelling sediments. Nature Geoscience, 3(8), 557–561. 10.1038/ngeo913

22. Gomes, F. M., Ramos, I. B., Wendt, C., Girard-Dias, W., De Souza, W., Machado, E. A., & Miranda, K. (2013). New insights into the in situ microscopic visualization and quantification of inorganic polyphosphate stores by 4’,6-diamidino-2-phenylindole (DAPI)-staining. European Journal of Histochemistry, 57(4), 228–236. 10.4081/ejh.2013.e34

23. Graue, J. (2007). Master thesis. Stickstoffixierung in einer marinen *Beggiatoa* Kultur. Universität Rostock, Rostock, Germany

24. Hansen, H. P., & Koroleff, F. (1999). Determination of nutrients. In K. Grasshoff, K. Kremling, & M. Ehrhardt (Eds.), Methods of Seawater Analysis (3rd ed., pp. 159–228). Wiley-VCH.

25. Himeoka, Y., Gummesson, B., Sørensen, M. A., Svenningsen, S. Lo, & Mitarai, N. (2022). Distinct Survival, Growth Lag, and rRNA Degradation Kinetics during Long-Term Starvation for Carbon or Phosphate. MSphere, 7(3). 10.1128/msphere.01006-21

26. Hinck, S., Mußmann, M., Salman, V., Neu, T. R., Lenk, S., de Beer, D., & Jonkers, H. M. (2011). Vacuolated *Beggiatoa*-like filaments from different hypersaline environments form a novel genus. Environmental Microbiology, 13(12), 3194– 3205. 10.1111/j.1462-2920.2011.02513.x

27. Hupfer, M., Gloess, S., & Grossart, H. (2007). Polyphosphate-accumulating microorganisms in aquatic sediment. Aquatic Microbial Ecology, 47, 299–311.

28. Jansson, M. (1993). Uptake, exchange, and excretion of orthophosphate in phosphate-starved *Scenedesmus quadricauda* and *Pseudomonas* K7. Limnol. Oceanogr, 38(6), 162–1178.

29. Jean, M. R. N., Gonzalez-Rizzo, S., Gauffre-Autelin, P., Lengger, S. K., Schouten, S., & Gros, O. (2015). Two new *Beggiatoa* species inhabiting marine mangrove sediments in the Caribbean. PLoS ONE, 10(2). 10.1371/journal.pone.0117832

30. Jørgensen, B. (1977). Distribution of Colorless Sulfur Bacteria (*Beggiatoa* spp.) in a Coastal Marine Sediment. Marine Biology, 41.

31. Jørgensen, B. B., Dunker, R., Grünke, S., & Røy, H. (2010). Filamentous sulfur bacteria, *Beggiatoa* spp., in arctic marine sediments (Svalbard, 79°N). FEMS Microbiology Ecology, 73(3), 500–513. 10.1111/j.1574-6941.2010.00918.x

32. Kamp, A., Røy, H., & Schulz-Vogt, H. N. (2008). Video-supported analysis of *Beggiatoa* filament growth, breakage, and movement. Microbial Ecology, 56(3), 484–491. 10.1007/s00248-008-9367-x

33. Konopka, A. E., Klemer, A. R., Walsby, A. E., & Ibelings, B. W. (1993). Effects of macronutrients upon buoyancy regulation by metalimnetic *Oscillatoria agardhii* in Deming Lake, Minnesota. In Journal of Plankton Research, 15(9). http://plankt.oxfordjournals.org/

34. Kornberg, A., Rao, N. N., & Ault-Riché, D. (1999). Inorganic Polyphosphate: A Molecule of Many Functions. Annual Review of Biochemistry, 68, 89–125.

35. Lake, B. A., Coolidge, K. M., Norton, S. A., & Amirbahman, A. (2007). Factors contributing to the internal loading of phosphorus from anoxic sediments in six Maine, USA, lakes. Science of the Total Environment, 373(2–3), 534–541. 10.1016/j.scitotenv.2006.12.021

36. Langer, S. (2019). PhD thesis. Polyphosphate in Marine Environments and *Beggiatoa* sp. – Quantification and Visualization. Universität Rostock, Rostock, Germany

37. Langer, S., Vogts, A., & Shuls-Vogt, H. (2018). Simultaneous Visualization of Enzymatic Activity in the Cytoplasm and at Polyphosphate Inclusions in *Beggiatoa* sp. Strain 35Flor Incubated with^18^O-Labeled Water. mSphere, 3(6).

38. Larkin, J. M., & Strohl, W. R. (1983). *Beggiatoa*, *Thiothrix*, and *Thioploca*. Annual Review of Microbiology, 37, 341–367. 10.1146/annurev.mi.37.100183.002013

39. Li, J., & Dittrich, M. (2019). Dynamic polyphosphate metabolism in cyanobacteria responding to phosphorus availability. Environmental Microbiology, 21(2), 572–583. 10.1111/1462-2920.14488

40. Lloyd, K. G., Albert, D. B., Biddle, J. F., Chanton, J. P., Pizarro, O., & Teske, A. (2010). Spatial structure and activity of sedimentary microbial communities underlying a *Beggiatoa* spp. mat in a Gulf of Mexico hydrocarbon seep. PLoS ONE, 5(1). 10.1371/journal.pone.0008738

41. Martín, J. F., & Liras, P. (2021). Molecular mechanisms of phosphate sensing, transport and signalling in streptomyces and related actinobacteria. International Journal of Molecular Sciences, 22(3), 1–20. 10.3390/ijms22031129

42. Martin, P., Dyhrman, S. T., Lomas, M. W., Poulton, N. J., & Van Mooy, B. A. S. (2014). Accumulation and enhanced cycling of polyphosphate by Sargasso Sea plankton in response to low phosphorus. Proceedings of the National Academy of Sciences of the United States of America, 111(22), 8089–8094. 10.1073/pnas.1321719111

43. McKay, L. J., MacGregor, B. J., Biddle, J. F., Albert, D. B., Mendlovitz, H. P., Hoer, D. R., Lipp, J. S., Lloyd, K. G., & Teske, A. P. (2012). Spatial heterogeneity and underlying geochemistry of phylogenetically diverse orange and white *Beggiatoa* mats in Guaymas Basin hydrothermal sediments. Deep-Sea Research Part I: Oceanographic Research Papers, 67, 21–31. 10.1016/j.dsr.2012.04.011

44. Medveczky, N., & Rosenberg, H. (1971). Phosphate Transport in *Escherichia Coli*. Biochimica et Biophysica Acta (BBA) - Biomembranes, 241(2), 494–506.

45. Möller, L., Laas, P., Rogge, A., Goetz, F., Bahlo, R., Leipe, T., & Labrenz, M. (2019). Sulfurimonas subgroup GD17 cells accumulate polyphosphate under fluctuating redox conditions in the Baltic Sea: possible implications for their ecology. ISME Journal, 13(2), 482–493. 10.1038/s41396-018-0267-x

46. Møller, M. M., Nielsen, L. P., & Jørgensen, B. B. (1985). Oxygen Responses and Mat Formation by *Beggiatoa* spp. Applied and Environmental Microbiology, 50(2), 373–382.

47. Murray, N. J., Phinn, S. R., DeWitt, M., Ferrari, R., Johnston, R., Lyons, M. B., Clinton, N., Thau, D., & Fuller, R. A. (2019). The global distribution and trajectory of tidal flats. Nature, 565(7738), 222–225. 10.1038/s41586-018-0805-8

48. Mußmann, M., Schulz, H. N., Strotmann, B., Kjær, T., Nielsen, L. P., Rosselló-Mora, R. A., Amann, R. I., & Jørgensen, B. B. (2003). Phylogeny and distribution of nitrate-storing *Beggiatoa* spp. in coastal marine sediments. Environmental Microbiology, 5(6), 523–533. 10.1046/j.1462-2920.2003.00440.x

49. Nelson, D. C., & Jannasch, H. W. (1983). Chemoautotrophic growth of a marine *Beggiatoa* in sulfide-gradient cultures. Archives of Microbiology, 136(4), 262–269. 10.1007/BF00425214

50. Noffke, A., Sommer, S., Dale, A. W., Hall, P. O. J., & Pfannkuche, O. (2016). Benthic nutrient fluxes in the Eastern Gotland Basin (Baltic Sea) with particular focus on microbial mat ecosystems. Journal of Marine Systems, 158, 1–12. 10.1016/j.jmarsys.2016.01.007

51. Ogle, H. D., Doll, C. J., Wheeler, A. P., & Dinno, A. (2023). FSA: Simple Fisheries Stock Assessment Methods (R package version 0.9.5). https://fishr-core-team.github.io/FSA/

52. Onofri, A. (2020). The broken bridge between biologists and statisticians: a blog and R package. Statforbiology. https://www.statforbiology.com

53. Pasek, M. A. (2008). Rethinking early Earth phosphorus geochemistry. The Proceedings of the National Academy of Sciences (PNAS), 105(3), 853–858. www.pnas.orgcgidoi10.1073pnas.0708205105

54. Pauli, A. S. L., & Kaitala, S. (1997). Phosphate uptake kinetics by Acinetobacter isolates. Biotechnology and Bioengineering, 53(3), 304–309. 10.1002/(SICI)1097-0290(19970205)53:3<304::AID-BIT9>3.0.CO;2-M

55. R core team. (2023). R: A Language and Environment for Statistical Computing (4.3.2). R Foundation for Statistical Computing. https://www.R-project.org/

56. Rao, N. N., Gómez-García, M. R., & Kornberg, A. (2009). Inorganic polyphosphate: Essential for growth and survival. In Annual Review of Biochemistry (Vol. 78, pp. 605–647). 10.1146/annurev.biochem.77.083007.093039

57. Rashid, M. H., & Kornberg, A. (2000). Inorganic polyphosphate is needed for swimming, swarming, and twitching motilities of *Pseudomonas aeruginosa*. The Proceedings of the National Academy of Sciences, 97(9), 4885–4890. www.pnas.org

58. Ravin, N. V., Rudenko, T. S., Beletsky, A. V., Smolyakov, D. D., Mardanov, A. V., Grabovich, M. Y., & Muntyan, M. S. (2024). Phylogeny and Metabolic Potential of New Giant Sulfur Bacteria of the Family Beggiatoaceae from Coastal-Marine Sulfur Mats of the White Sea. International Journal of Molecular Sciences, 25(11). 10.3390/ijms25116028

59. Riegman, R., & Mur, L. R. (1984). Phosphate uptake by P-limited Oscillatoria agardhii. FEMS Microbiology Letters, 21(3), 335–339. 10.1111/j.1574-6968.1984.tb00332.x

60. Romano, S., Schulz-Vogt, H. N., González, J. M., & Bondarev, V. (2015). Phosphate limitation induces drastic physiological changes, virulence-related gene expression, and secondary metabolite production in *Pseudovibrio* sp. strain FO-BEG1. Applied and Environmental Microbiology, 81(10), 3518–3528. 10.1128/AEM.04167-14

61. Saia, S. M., Carrick, H. J., Buda, A. R., Regan, J. M., & Todd Walter, M. (2021). Critical review of polyphosphate and polyphosphate accumulating organisms for agricultural water quality management. In Environmental Science and Technology, 55(5), pp. 2722–2742. American Chemical Society. 10.1021/acs.est.0c03566

62. Santoro, M., Hassenrück, C., Labrenz, M., & Hagemann, M. (2023). Acclimation of *Nodularia spumigena* CCY9414 to inorganic phosphate limitation – Identification of the P-limitation stimulon via RNA-seq. Frontiers in Microbiology, 13. 10.3389/fmicb.2022.1082763

63. Schulz, H. N., & Schulz, H. D. (2005). Large sulfur bacteria and the formation of phosphorite. Science, 307(5708), 416–418. 10.1126/science.1103096

64. Schwedt, A. (2011). PhD Thesis. Physiology of a marine *Beggiatoa* strain and the accompanying organism *Pseudovibrio* sp. – a facultatively oligotrophic bacterium., Rostock University, Rostock, Germany.

65. Seeger, M., & Jerez, C. A. (1993). Phosphate-starvation induced changes in Thiobacillus ferrooxidans. FEMS Microbiology Letters, 108(1), 35–41. 10.1111/j.1574-6968.1993.tb06070.x

66. Shiba, T., Tsutsumi, K., Yano, H., Ihara, Y., Kameda, A., Tanaka, K., Takahashi, H., Munekata, M., Rao, N. N., & Kornberg, A. (1997). Inorganic polyphosphate and the induction of rpoS expression. Biochemistry, 94, 11210–11215. www.pnas.org.

67. Solovchenko, A., Gorelova, O., Karpova, O., Selyakh, I., Semenova, L., Chivkunova, O., Baulina, O., Vinogradova, E., Pugacheva, T., Scherbakov, P., Vasilieva, S., Lukyanov, A., & Lobakova, E. (2020). Phosphorus feast and famine in cyanobacteria: Is luxury uptake of the nutrient just a consequence of acclimation to its shortage? Cells, 9(9), 1–21. 10.3390/cells9091933

68. Strohl And, W. R., & Larkin, J. M. (1978). Enumeration, Isolation, and Characterization of *Beggiatoa* from Freshwater Sediments. Applied and Environmental Microbiology, 36(5), 755–770. https://journals.asm.org/journal/aem

69. Van Veen, H. W., Abee, T., Kortstee, G. J. J., Konings, W. N., & Zehnder14, A. J. B. (1993). Characterization of Two Phosphate Transport Systems in *Acinetobacter johnsonii* 210A. In Journal of Bacteriology. https://journals.asm.org/journal/jb

70. Werner, T. P., Amrhein, N., & Freimoser, F. M. (2005). Novel method for the quantification of inorganic polyphosphate (iPoP) in Saccharomyces cerevisiae shows dependence of iPoP content on the growth phase. Archives of Microbiology, 184(2), 129–136. 10.1007/s00203-005-0031-2

